# Multiscale Mechanics of Granular Biofilms

**DOI:** 10.1101/2025.06.18.660425

**Authors:** Lisa K. Månsson, Anna E. Warsaw, Elizabeth G. Wilbanks, Angela A. Pitenis

## Abstract

Biofilms produce and maintain extracellular polymeric substances (EPS) essential for their form and function. While biofilms are commonly lamellar and frequently targets of removal, granular biofilms are increasingly incorporated into water treatment strategies. In both cases, the EPS (mainly consisting of proteins, polysaccharides, and extracellular DNA) is largely responsible for their persistence. Unlike many granular biofilms, which are formed in engineered industrial bioreactors, the “pink berry” consortia is a naturally-occurring and robust granular biofilm of photosynthetic bacteria, found only in intertidal pools of salt marshes around Woods Hole, Massachusetts (USA). The pink berry biofilm’s unique ecological niche has sparked over three decades of study, yet their mechanical properties are completely unknown. Here, we characterized the structural and mechanical landscape of pink berry granules to determine the extent to which microscale heterogeneity influences macroscale material properties. We performed microindentation measurements on intact granules and nanoindentation measurements on thin sections. We report that intact pink berry granules exhibited low reduced elastic moduli (*E\**_pink berry_ ≈ 0.5–10 kPa) and fast stress relaxation times (τ_1/2_ ≈ seconds), consistent with previous investigations of soft and viscoelastic biofilms. Nanomechanical measurements of thin pink berry sections revealed two mechanically-distinct domains: a very soft extracellular polymeric substance (EPS) matrix surrounding stiffer microcolonies of purple sulfur bacteria (PSB). Light sheet fluorescence microscopy revealed the spatial organization and distribution of cell-dense PSB microcolonies (34 vol.%) within EPS matrix (66 vol.%), suggesting the nanomechanical behavior of EPS dominates macroscale pink berry mechanics. Our multiscale experimental approach combining mechanics and imaging may be broadly applicable to investigations of complex soft materials, from synthetic hydrogel composites to biologically heterogeneous spheroids, organoids, and tissues.

## 1. Introduction

Multicellularity critically depends on the ability to stick together under diverse stressors. Many multicellular systems, from bacterial aggregates to organs, generate an extracellular matrix as a protective binder and conduit for nutrient transport. Light and electron microscopy reveal that the extracellular matrix supports intricate structural complexity, spanning the macro- to nano-scales [1–3]. Nanomechanical measurements of single cells (e.g., atomic force microscopy) provide mechanobiological insights at the cellular and subcellular scales [4–12], yet large multicellular assemblies often exhibit very different force responses to mechanical compression due to spatial heterogeneity [13]. Such mechanical differences from the cellular scale to macroscale capture key emergent properties of multicellular systems – phenotypes that are foundational in the evolution of group-level traits from cellular-level activities [14].

Among the most ancient multicellular assemblies are communities of microorganisms held together by a self-produced extracellular matrix, also known as biofilms [15]. Mechanical measurements of biofilms have typically focused on lamellar biofilms relevant to both biomedical and industrial environments [10–12, 16–27]. However, millimeter-sized roughly spherical “granular” biofilms are observed in both natural and engineered environments and play a critical role in industrial wastewater treatment technologies [28–31]. Dense granules settle quickly, which serves to retain catalytically-valuable biomass within wastewater treatment reactors, and rapid respiration rates depletes oxygen at the granule core, leading to biofilms that can simultaneously catalyze both aerobic and anaerobic processes [28–31]. One of the few well-studied, naturally occurring granular biofilms are the colorfully described “pink berry” microbial consortia [32]. Pink berries are pink-hued granules, millimeters in diameter, found in intertidal salt marsh ponds (**Fig. 1**). Pink berries have enjoyed a long history of observation, leading to a genome-resolved understanding of the microbial community within the consortia [32–36]. Our prior work has established that these granules host a conserved consortia of bacteria (and diatoms), and are primarily composed of a single species of uncultivated phototrophic purple sulfur bacteria (*Thiohalocapsa* PSB1), which forms dense microcolonies within a translucent, supportive matrix of extracellular polymeric substances (EPS). Despite their importance to pink berries’ remarkable robustness and persistence in nature [32] and potential for industrial applications, their mechanical properties are completely unknown.

**Figure 1:**
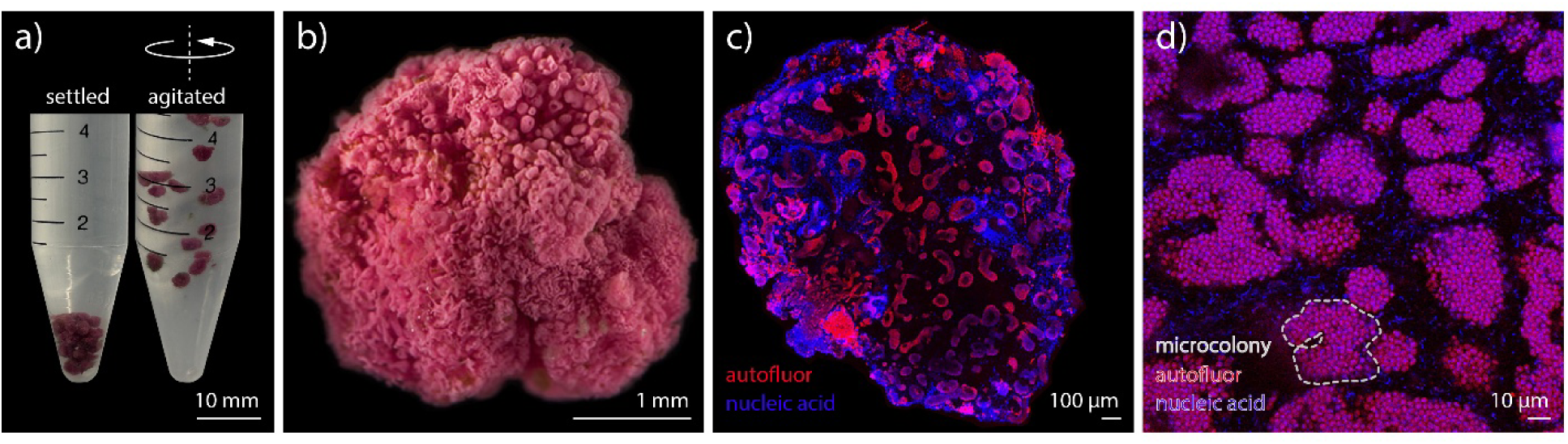
a) Pink berries settle quickly (left) and maintain integrity during agitation (right). b) Close-up view of a pink berry granular biofilm, reproduced from reference [35]. c) Fluorescence micrograph of a thin pink berry slice, with nucleic acid in blue (SYTO 41 Blue Fluorescent Nucleic Acid Stain) and cyanobacteria and purple sulfur bacteria (PSB) microcolony autofluorescence in red. d) Higher magnification view of PSB microcolonies (encircled in dashed line) within a pink berry slice, with nucleic acid in blue (DAPI) and PSB microcolony autofluorescence in red.

In this study, we present multiscale structure-property relationships of pink berry granular biofilms. We conducted in situ micromechanical measurements on intact granules as well as in situ nanomechanical measurements on thin sections to distinguish the respective contributions of bacterial microcolonies and the EPS matrix to the mechanics of the granular composite. We then leveraged advanced imaging approaches to correlate granule structure and spatial organization with quantitative measurements of their mechanical properties.

## 2. Materials and Methods

### 2.1. Sample Collection

Pink berry granular biofilms were sampled in September 2024 from an intertidal pond in the Little Sippewissett Salt Marsh in Falmouth, MA (N 41.57587, W -70.63923). Pink berries were collected with a sieve (1 mm mesh size) from the sediment-water interface, stored in water sampled from the site, and shipped to UC Santa Barbara. The pink berries were maintained as a concentrated collection in an aquarium with ≈3 cm sediment sampled from the site and ≈6 L unfiltered seawater.

### 2.2. Microindentation of Pink Berries

Pink berries were collected from the aquarium (never frozen) and washed three times in 0.2 µm filter-sterilized seawater. The granular biofilms were fully submerged in phosphate-buffered saline (PBS 3×) and positioned in a rigid transparent dish for unconfined compression using a custom-built microtribometer [37–43]. Pink berry deformation and recovery were recorded using an inverted digital microscope (Dino-Lite Edge AF4115ZT, 20–220× magnification) mounted beneath the dish. Pink berries were compressed at a constant approach and retraction velocity (*v* = 1, 10, or 100 µm/s) to a maximum applied normal force between *F*_N_ = 500 µN and 1 mN (contact pressures, *P* = 80 Pa to 330 Pa) using a hemispherical glass probe (radius of curvature, *R*_1_ = 2.6 mm) attached to a double-leaf cantilever (normal spring stiffness, *K*_N_ = 220 µN/µm). Each pink berry sample was indented at least three consecutive times (n = 3) with a minute between each indent to obtain the average elastic modulus.

For calculation of reduced elastic modulus, *E*\*, the data were fit to the Hertzian contact mechanics model [44] using Eq. 1.

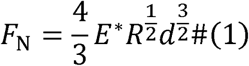

In this equation, *F*_N_ is the applied normal force, *d* is the microindentation depth, and *R* is the effective radius determined from the probe radius of curvature, *R*_1_, and the radius of the pink berry sample, approximated as a sphere, *R*_2_ (Eq. 2). The average pink berry diameter for each sample was determined from five measurements across each sample (**Supplementary Fig. A1**).

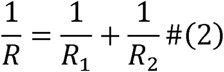

To comply with Hertzian contact mechanics model assumptions, all indentation measurements were analyzed within the small-strain regime (i.e., maximum contact radius *a* < 0.15*R*_2_) (**Supplementary Fig. A2**).

### 2.3. Stress Relaxation Measurements

Pink berry samples were washed and positioned on a rigid dish for unconfined compression while fully submerged in PBS (3×) as described in Section 2.2. Using a cylindrical glass rod (radius, *R*_cyl_ = 0.75 mm), pink berry samples (N = 6) were compressed to an approximate strain of 5% at an approach velocity of *v*_app_ = 2 mm/s (strain rate ≍ 20 s^-1^), held for 20 s, then retracted at *v*_ret_ = 0.05 mm/s (motion profile in **Fig. 3b**). The data was smoothed using a Savitzky-Golay filter in MATLAB, the stress was normalized to the maximum stress, and the characteristic stress relaxation time constant, τ_1/2_, was determined at the time the stress had decreased to half of its maximum value. The average pink berry diameter for each sample was measured from images taken from below by the inverted digital microscope (**Supplementary Fig. A3**). For each sample, five diameters across were measured to obtain an average.

Stress relaxation data were fit to common viscoelastic models based on Maxwell elements where viscous contributions are represented by dashpots and elastic contributions by springs. The equations used are presented in Supplementary Information, with data fits in **Supplementary Fig. C1**.

### 2.3. Nanoindentation of Pink Berry Slices

To investigate the mechanical properties of distinct biofilm components, pink berries were prepared for sectioning. A cylindrical disk of agarose was prepared by pouring 4% (w/v) agarose gel in PBS (3×) into a rigid mold (≈20 mm diameter, ≈10 mm thickness). Prior to gelation of agarose at room temperature, a single pink berry was embedded live (unfixed) in the center of the mold. Thereafter, the cylindrical disk was sectioned on a Leica VT1000 S vibratome to obtain roughly 100 µm thick slices through the entire disk. Pink berry slices were stored in PBS (3×) in glass-bottom culture dishes at 4°C for up to one week prior to testing. During this time, negligible sample degradation was observed. To prepare slices for nanoindentation, a glass-bottom culture dish was coated with 400 µL of Weaver’s Gelatin (50 mL DI H_2_O, 0.5 g chromium potassium sulfate dodecahydrate, 0.05 g gelatin type B) and incubated for 10 min at room temperature. The solution was aspirated, and the glass surface dried for 15 min at room temperature. Pink berry slices were gently patted with cellulose paper to remove excess liquid, positioned on the coated glass surface, and equilibrated for 20 min at room temperature. The culture dish was then filled with PBS (3×) for nanoindentation measurements. The full procedure is illustrated in **Supplementary Fig. B1**.

Nanoindentation measurements were performed with the Optics11 Life Pavone housed in the BioPACIFIC MIP user facilities at UC Santa Barbara. Pink berry slices were indented at room temperature using a spherical glass probe (probe radius of curvature, *R* = 26.5 µm, normal spring constant, *K*_N_ = 0.021 N/m) while fully submerged in PBS (3×) at a constant indentation velocity of *v* = 10 µm/s to an indentation depth of about 5 µm. Multiple positions across each slice (separated by at least two probe diameters) were selected using the inverted 20× objective installed in the instrument. Each position across the pink berry slice was indented at least three consecutive times in the same location to obtain an average elastic modulus. All measurements were conducted at least two probe diameters from the perimeter of the pink berry, which is near the interface between the biofilm and the agarose gel matrix. The mechanical properties of the agarose support were characterized following the procedures above using a probe with appropriate cantilever spring constant for the material (probe radius of curvature, *R* = 48 µm, normal force spring constant, *K*_N_ = 3.52 N/m) (agarose elastic modulus was about 500 kPa, see **Supplementary Fig. B2**, in agreement with previous literature [45]). Following nanoindentation measurements, the full pink berry slices were imaged to correlate micro- to -macro-scale structure. Slices were imaged using an inverted Leica SP8 confocal microscope. A 10× (0.3 NA) air objective lens was combined with tile scanning to image entire pink berry slices. Brightfield images were acquired using 3×3 tiles with 10% overlap.

The reduced elastic modulus (*E*\*) was calculated for nanoindentation measurements as described above, with *R* in Eq. 1 here being the radius of the spherical probe. Statistical analyses were performed in Python using the *scipy* package, specifically the function *mannwhitneyu* for performing a Mann-Whitney u-test, as the nanoindentation data from purple sulfur bacteria (PSB) microcolonies were not normally distributed.

### 2.5. Fluorescence Light Sheet Microscopy and Analysis

To examine the internal architecture of the pink berry granular biofilms, a pink berry collected from an intertidal pond in the Great Sippewissett Salt Marsh, Falmouth, MA (N 41.58618, W - 70.64228) was fixed, optically cleared, and imaged on a light sheet microscope. Briefly, the pink berry was twice washed in 0.2 µm-filtered seawater, fixed in 4% PFA in PBS (1×) for 1 h agitated at 125 rpm at room temperature, washed three times in PBS (1×), and stored in PBS (1×) at 4 °C for about one week. The pink berry was incubated in 100% methanol for about 2 h, twice washed in PBS (1×), and incubated in 500 µL clearing reagent iCBiofilm-H2 (TCI AMERICA) diluted 4:1 with sterile, de-ionized water for about 7 h. For imaging, the pink berry was embedded in 1% (w/v) agarose gel in the 4:1 iCBiofilm-H2:water solution. The agarose gelled within the barrel of a 1 mL plastic disposable syringe modified with a wider inlet to reduce compressive strains on the pink berry during aspiration. The syringe was stored at 4 °C in the 4:1 iCBiofilm-H2:water solution until imaging.

Light sheet imaging was performed using the Zeiss Lightsheet Z.1 microscope housed in the NRI-MCDB shared facility using the 20× CLARITY objective lens (refractive index = 1.46) combined with tile scanning. The sample chamber was filled with iCBiofilm-H2 solution, and the pink berry embedded in agarose was partially extruded from the syringe and immersed in the solution. We acquired a Z-stack through 844 µm of the pink berry (670 slices, Z-step = 1.26 µm), spanning roughly half of the granule, and 3×4 tiles were imaged with 10% overlap. A 488 nm laser at 10% intensity was used to excite pink berry autofluorescence, and emission was collected at 505–545 nm.. The tiles were stitched together in Fiji (ImageJ) using the stitching plugin Grid/Collection Stitching [46]. Image analysis was conducted using Imaris 10.2.0. Three surface categories were created from the Z-stack, capturing the intact pink berry surface, bright abiotic crystals, and PSB microcolonies. These surfaces were created using the machine learning segmentation tool in Imaris, which trains an AI segmentation model. One of the surface categories captured bright crystals that appeared in the Z-stack as an artifact of the clearing protocol and was used to remove these from the images. The microcolony category primarily captured PSB microcolonies but also included a small number of potential cyanobacterial and *Rhodobacteraceae* microcolonies. The volumes computed from the pink berry and microcolony surfaces were used to calculate the microcolony volume fraction.

## 3. Results

### 3.1. Pink Berry Granular Biofilms are Soft and Exhibit Low Elastic Modulus

Microindentation measurements of whole pink berries (i.e., granules) were performed with a custom-built microindentation platform using the experimental configuration shown in **Fig. 2a**. A representative force-displacement approach curve for a constant indentation velocity of *v* = 10 µm/s is shown in **Fig. 2b**. Granules exhibited a range of reduced elastic moduli, from 0.5 < *E*\*_pink berry_ < 11.2 kPa, with an average of *E*\*_pink berry_ = 4.0 ± 3.4 kPa (N = 7). Although there was no clear correlation with pink berry diameter (**Fig. 2c**), the wide range of elastic modulus values obtained in this study is consistent with previously observed heterogeneity of biofilms, even in pure culture of single-species systems [47]. The reduced moduli of granules are within comparable order of magnitude to lamellar biofilms (see additional discussion in section 4.1) [17]. Serial force-displacement curves on the same location of the same granule did not reveal significant adhesion, plastic deformation, or strain-stiffening (**Supplementary Fig. A4)**. The reduced elastic modulus of pink berry granules was largely insensitive to indentation velocity (*v* = 1, 10, and 100 µm/s) (**Supplementary Fig. A5**).

**Figure 2:**
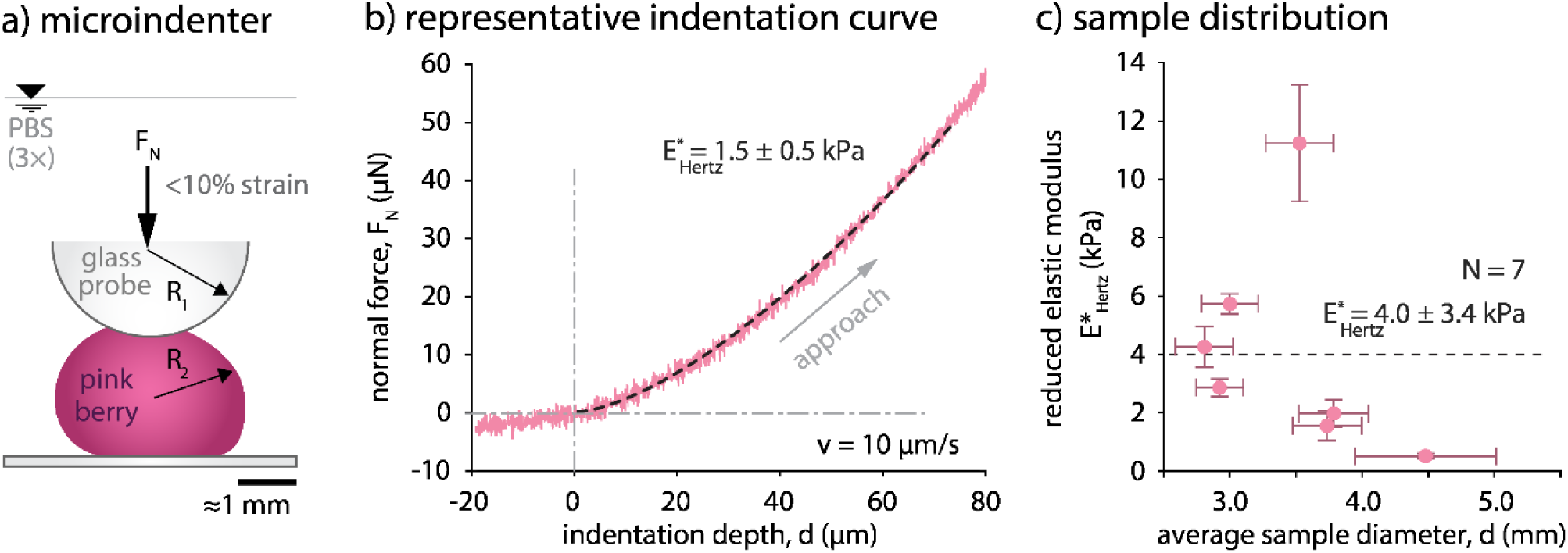
Microindentation measurements of whole pink berries. a) Experimental configuration for microindentation measurements. b) Representative force-displacement curve for a single pink berry sample with approach and retraction paths indicated. The Hertzian contact mechanics model (dashed line) was used to fit the approach curve to determine the elastic modulus. c) Pink berry granules reduced elastic modulus average values are shown with y-error bars indicating standard deviation from n = 3 indentations per sample, and x-error bars representing the standard deviation from the average sample diameter (n = 5).

### 3.2. Pink Berry Granular Biofilms Show Fast Response to Compression during Stress Relaxation

The custom-built microindenter with a modified experimental configuration (flat punch glass probe) was used to conduct stress relaxation measurements on individual pink berry granules (**Fig. 3a,b**). Pink berry samples were compressed to an average strain of 5% (N = 6). Normalized stress relaxation data was used to determine the characteristic relaxation time, τ_1/2_, the time required for the applied stress to reach half its initial value (**Fig. 3c**). Stress relaxation time was on the order of 0.1 to 10 s, with an average stress relaxation time τ_1/2_ = 3.7 ± 4.1 s (N = 6) (**Fig. 3c**). Stress relaxation time for the small sample size evaluated here does not appear to be correlated with sample diameter (**Fig. 3d**). Stress relaxation studies performed by compression testing of dental biofilm (plaque) report similar time scales, τ_1/2_ = 11 ± 12 s (n = 10), and similar variability, attributed to heterogeneity associated with biofilms [47]. Collagen biopolymers and heart tissues under modestly higher strains (10–15%) exhibit similar relaxation times extracted from indentation measurements [48, 49]. Notably, decellularized heart tissue was found to relax stress slower than the heart tissue containing cells [49]. This finding suggests an intriguing possibility that the cell microcolony component in the pink berry may make important contributions to the granule’s fast response to compression. Further studies could include stress relaxation measurements of decellularized samples to examine this hypothesis.

**Figure 3:**
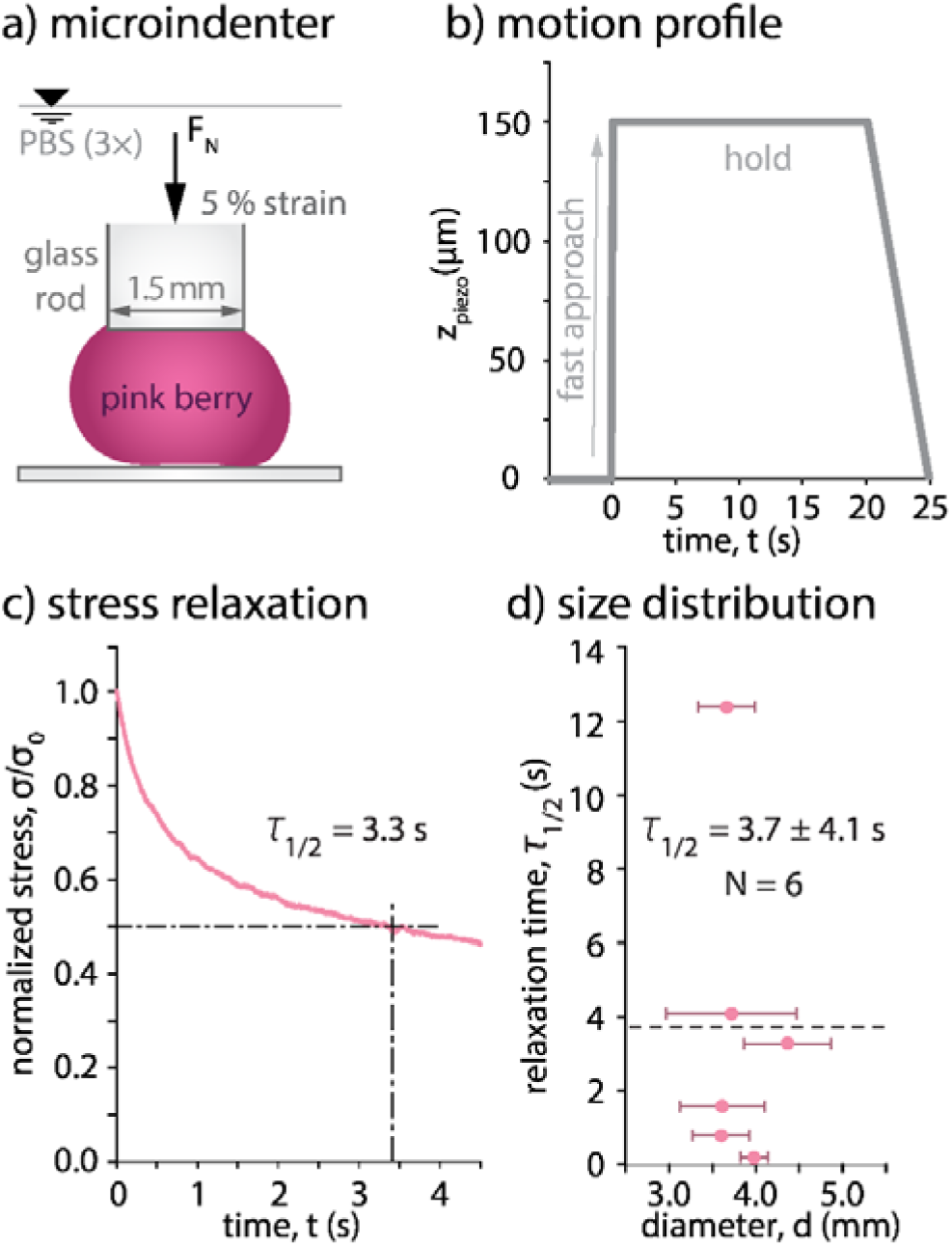
Pink berry granular biofilms exhibit fast relaxation during compression. a) Experimental configuration of stress relaxation measurements. b) Microindenter motion profile. c) Resulting stress relaxation curve from microindentation using the motion profile in b), where the stress was normalized to the maximum stress indented. Each pink berry was evaluated only once (N = 6, n = 1). d) Stress relaxation time scale shows no dependence on pink berry sample size. Error bars in x show standard deviations for each sample’s average diameter.

### 3.4. Nanoindentations Reveal Microscopic Mechanical Heterogeneity in Pink Berries

Pink berry biofilms are composed of tightly packed clusters of coccoid purple sulfur bacterial cells (PSB microcolonies) supported within an EPS matrix (**Fig. 1**). To separately measure these distinct structures and examine their contributions to the mechanical properties of the whole granule, we sectioned pink berry samples (about 100 µm thin) for nanoindentation measurements on either microcolonies or EPS domains (**Fig. 4a**). Nanoindentation measurements were conducted using the Optics11 Life Pavone nanoindenter with an integrated inverted 20× objective and a borosilicate glass probe (*R* = 26.5 µm radius). Imaging through thin sections of pink berry granules enabled unobstructed observations of purple sulfur bacteria (PSB) microcolonies surrounded by EPS (**Fig. 4b**), with the location of each nanoindentation measurement represented by circles (red for microcolonies and blue for EPS). The apparent contact diameter, 2*a*, is represented to scale in the inset (**Fig. 4b**), and indentations were separated by a minimum distance of 100 µm from neighboring locations. Additionally, each location was selected such that the projected contact area, as observed by inverted microscope, was fully bounded by either the microcolony or EPS domains to reduce crosstalk.

**Figure 4:**
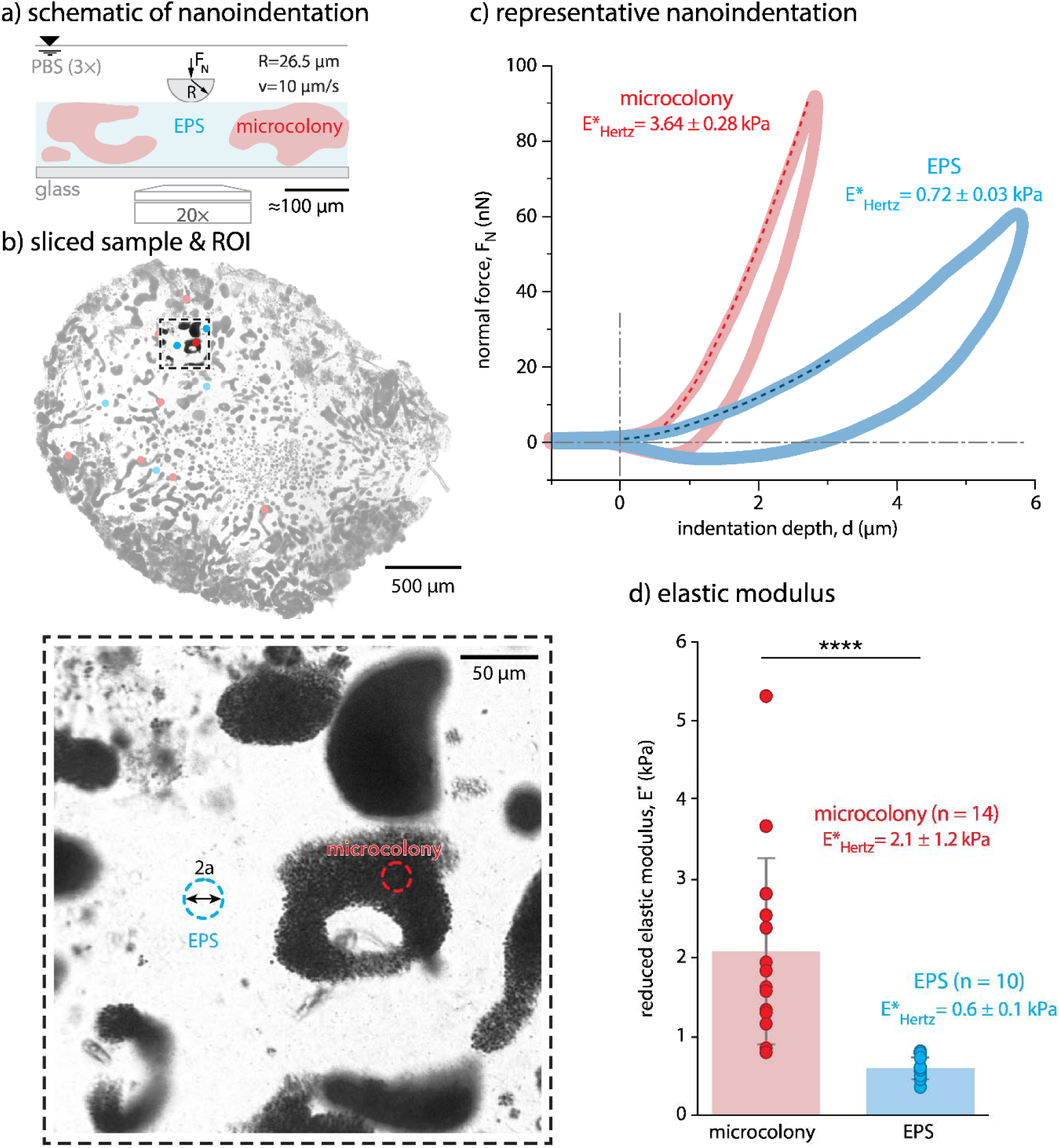
Nanoindentations of sliced pink berries reveal heterogeneity in mechanical properties at the micrometer scale. a) Schematic side view of experimental configuration. b) Brightfield image of the pink berry section, showing purple sulfur bacteria (PSB) (dark) and extracellular polymeric substances (EPS) (light) and probe contact radius, *a*, determined at maximum indentation depth. c) Representative force-displacement curves from nanoindentation measurements in positions shown in b). Hertzian contact mechanics model (dashed lines) used to fit approach curves. d) PSB microcolonies exhibited significantly higher elastic moduli than EPS matrix.

Nanoindentation measurements were conducted over two slices from a single pink berry granule (N = 2). Image maps of pink berry slices with indentation locations marked along with corresponding reduced elastic modulus values are presented in **Supplementary Fig. B3**.

Nanoindentation measurements revealed significant differences in reduced elastic modulus between PSB microcolonies (*E\**_PSB_ = 2.1 ± 1.2 kPa, n = 14 locations, each location with 3 technical repeats) and the EPS (*E\**_EPS_ = 0.6 ± 0.1 kPa, n = 10 locations, each location with 3 technical repeats) (**Fig. 4c,d**). The average reduced elastic modulus between the PSB microcolonies and EPS matrix domains were determined to be statistically significant based on the Mann Whitney test (p-value = 0.000002). The reduced elastic modulus determined from nanoindentation measurements against PSB microcolonies ranged from *E*\*_PSB_ = 0.8 to 5.3 kPa, and from *E*\*_EPS_ = 0.4 to 0.8 kPa for the EPS matrix. The reported elastic modulus of the EPS matrix within pink berry granules is on the same order of magnitude as those determined from the EPS found in lamellar biofilms [50]. The broad spread of reported *E\** values associated with PSB microcolonies (**Fig. 4** and **Supplementary Fig. B3**) may be due to a number of factors, including: biofilm heterogeneity, microcolony size and density, and depth into the pink berry slice. Nanoindentation locations were selected using optical microscopy; opaque PSB domains may have occluded EPS matrix above the microcolony and directly beneath the probe, which may have led to unintended crosstalk.

### 3.5. Volume Fraction of Microcolonies Distributed Throughout the EPS Matrix

To examine the internal structure of pink berry biofilms in 3D, we used fluorescence light sheet microscopy commonly used in neurobiology and developmental biology to image organs and embryos [51]. This technique images millimeter-sized samples with fast data acquisition, using tissue clearing and purely optical sectioning [52] and allows us to capture the internal architecture of an intact, millimeter-scale pink berry biofilm with single cell resolution (**Fig 5a**). Imaging revealed heterogenous, irregularly-shaped microcolonies distributed throughout the granule (**Fig. 5a**). The EPS matrix between microcolonies featured occasional diatoms. A cropped region of the granule shows the microcolonies in greater detail (**Fig. 5b**), which can be described as roughly tube shaped, about 30 µm in diameter on average, often curved and sometimes connected into a toroidal-like shape (**Fig. 5b,c**). Representative Z-slices through the 3D image show the spatial organization of PSB microcolonies as well as occasional bright masses of diatoms or cyanobacteria (e.g., in the upper center part of the second to last slice shown in **Fig. 5d**).

**Figure 5:**
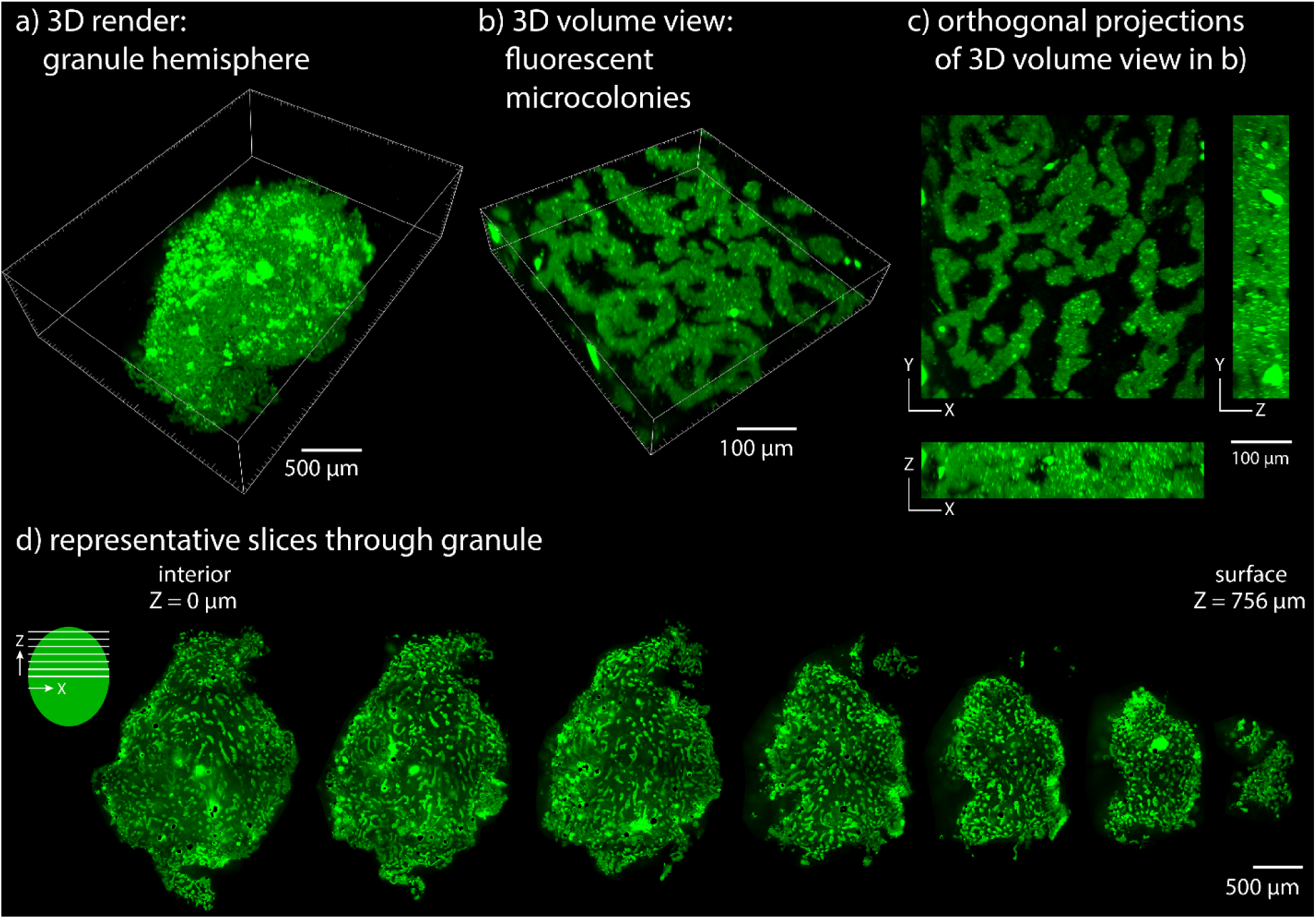
Light sheet microscopy reveals heterogeneous microcolonies distributed throughout the pink berry granular biofilm. a) Full 3D rendering of a Z-stack through half a pink berry granule. b) Cropped region of 3D rendering (500 × 500 × 100 µm) showing the structure of PSB microcolonies in green (autofluorescence). c) Orthogonal views of cropped region shown in b) in the XY, XZ, and YZ directions. d) Representative Z slices through Z-stack shown in a), from Z = 0 µm at the interior of the pink berry granule to Z = 756 µm near the surface. Each slice shown is 126 µm apart.

To further analyze pink berry composition, we used machine learning to obtain volumetric surface renderings capturing different components of the pink berry granular biofilm (**Fig. 6**). Using a surface reconstruction of the granule, we determined the volume to be 1.86 mm^3^ (**Fig. 6a, 6b**). Surface reconstruction of each microcolony gave a summed microcolony volume of 0.629 mm^3^ (**Fig. 6c, 6d**). In this granule, the volume fraction of microcolonies corresponded to *f*_PSB_ ≈34% of the pink berry volume and the EPS matrix made up a volume fraction of *f*_EPS_ ≈66%.

**Figure 6:**
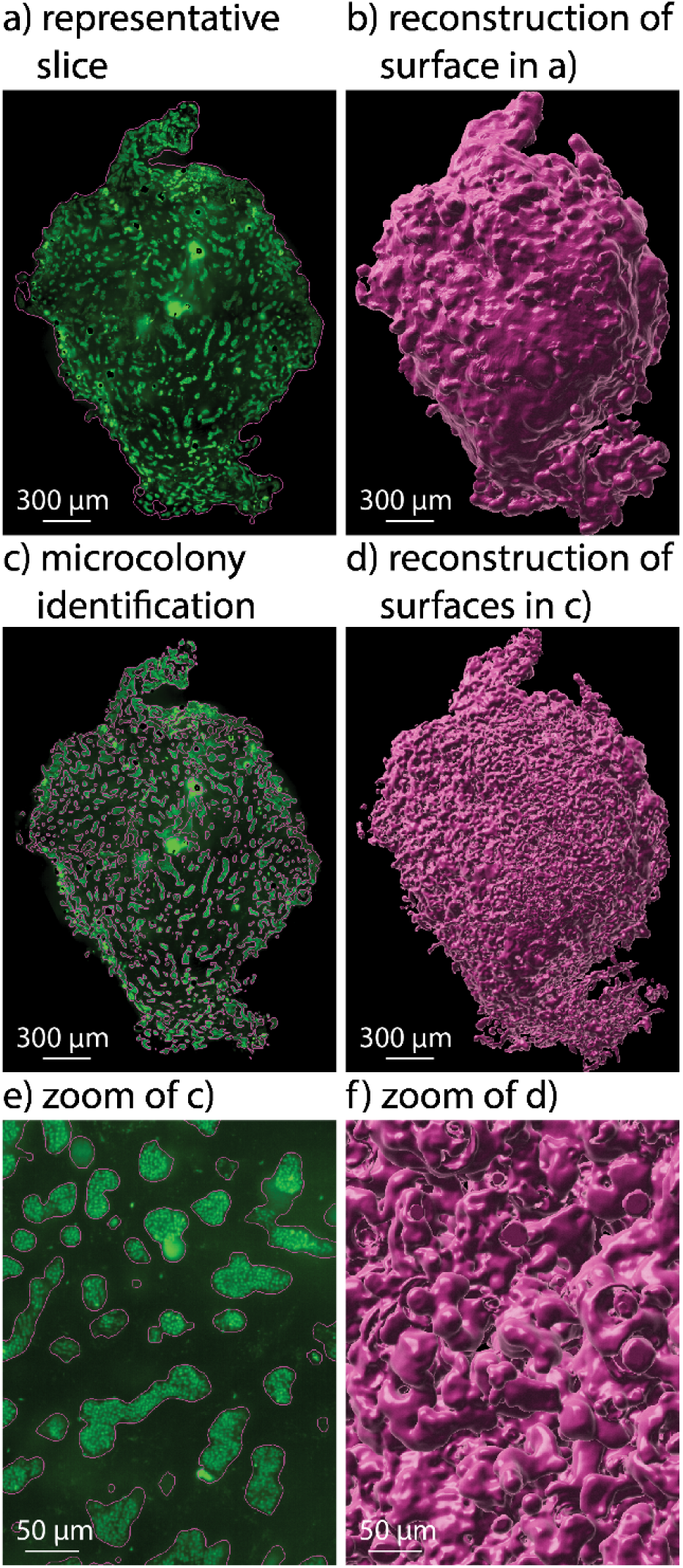
Microcolonies correspond to roughly 34% of the entire pink berry volume. a) Representative slice showing autofluorescent microcolonies (green) and the outline in pink representing the granule surface reconstruction shown in b. b) Granule surface reconstruction from a Z-stack through half a pink berry granule. Imaris was used to obtain the surface via machine learning segmentation. c) Representative slice illustrating autofluorescent microcolonies (green) and circled in pink is the microcolony surface rendering shown in d. d) Microcolony surface reconstruction from the same Z-stack using Imaris machine learning segmentation. e) Enlarged view of c to showcase microcolonies. f) Enlarged view of d.

## 4. Discussion

### 4.1 Connecting Structure to Mechanical Properties Across Scales

In this study, we present methods that bridge multiscale mechanical properties and the structural landscape of composite biological samples. We performed mechanical measurements using indentation techniques across nN to µN force scales. Advanced imaging using light sheet fluorescence microscopy allowed us to visualize intact macroscopic specimens with micron-scale resolution, connecting our smaller scale, high-resolution imaging methods and larger-scale visualization of entire macroscopic specimens [53]. Light sheet fluorescence microscopy connected structural visualizations to the same length scales as our mechanical characterization. Previously, to resolve single-celled structures in this system, we relied on physical thin sectioning of an embedded sample, combined with confocal microscopy [33]; given the technical challenges of capturing and aligning serial thin sectioning, this approach was ill-suited to reconstructing finely sampled millimeter-sized structures. Here, we demonstrate a method that can reliably and rapidly (∼2 days) generate single-cell resolution images of intact, millimeter-sized granules. While light sheet imaging has occasionally been used to visualize surface-attached biofilms [54–58], this approach is not widespread in microbiology [59]. Our study is the first of its kind to use this approach to reveal the detailed cellular architecture of millimeter scale granular biofilms, and only the second reported application of this technique on marine bacteria [60].

We report reduced elastic moduli of whole pink berry granules in the range of *E*\*_pink berry_ ≈ 0.5–10 kPa, values within the many orders of magnitude wide range of elastic moduli values reported for lamellar biofilms, reviewed by Böl *et al.* [17]. However, it shall be noted that the elastic modulus determined depends on several factors including, for example, instrumentation used, specific experimental setup, biofilm species, and environmental conditions. Focusing only on measurements performed using microcantilever indentation or compression (between which, for example, probe geometries, indentation velocities, and sample strains still varied), elastic moduli for different lamellar biofilms (single-species, and with different cell-cell architecture) ranged from 0.017 kPa [61] up to 2.2 kPa [62]. The range of elastic moduli determined for pink berries is similar, although notably on the stiffer side. It should be noted that the Hertz contact mechanics model used to analyze microindentation measurements assumes indentation of a spherical sample (reflected in Eqs. 1 and 2) which can influence the reduced elastic modulus. The equations used imply that for an underestimated sample radius (*R*_2_ in Eq. 2), which may be the case for more oval-shaped pink berry samples, the estimated reduced elastic modulus *E*\* (from Eq.1), would be slightly overestimated.

This research effort attempted to connect the mechanical properties of whole, intact pink berries to their primary microscale constituents: PSB microcolonies and the EPS matrix. One pink berry among the samples characterized with microindentation was sectioned and further investigated via nanoindentation measurements. The largest granule (pictured in **Supplementary Fig. A1**, sample #5) was chosen for easier experimental handling. This sample was also the softest among the pink berries examined, with an average reduced elastic modulus of *E\**_pink berry_ = 0.5 ± 0.1 kPa (**Fig. 2c**). Bacterial microcolonies, estimated to comprise about 34 vol.% of the granule, exhibited an average reduced elastic modulus of *E\**_PSB_ = 2.1 ± 1.2 kPa. The predominant component, EPS (about 66 vol.% of the granule) was significantly softer, with an average reduced elastic modulus of *E\**_EPS_ = 0.6 ± 0.1 kPa. Our results suggest that the EPS dominates the overall mechanical response of the pink berry. Interestingly, similar EPS-dominant macroscopic mechanical properties have been previously reported for distinctly different lamellar biofilms [63]. The composition of EPS, which has been well characterized in several lamellar biofilms and includes a suite of polysaccharides, extracellular proteins and extracellular DNA, largely consists of water (around 95 to 98% water [64]) which consequently drives mechanical and transport properties of biofilms [65]. We hypothesize that softer EPS enables greater accommodation of deformation, thus protecting microcolonies from cell damage and preventing intra-granular fracture, even under moderate compressive strains (about 50% strain, by manual inspection).

EPS is an essential component of granular biofilms that enables granules to hold together under flow in bioreactors [31]. The EPS in naturally occurring pink berry biofilms has been modified by nature over time, and our results demonstrate that the EPS matrix is a key component in these highly resilient structures. Pink berry EPS and the species that produce it may be useful in optimizing granules for industrial bioreactors, or for the rational design of engineered microbial consortia, an ongoing challenge in synthetic biology [66–68]. Ultimately, this may be useful both in engineering robust and functional industrial systems, as well as inspiring novel 3D-structured soft materials.

### 4.2 Pink Berry Permeability

Microcolonies of cells within the pink berry require nutrient supply through fluid flow. Pink berry granules are estimated to contain about 70% water; the interconnected domains of EPS may provide channels for fluid flow through the granule [17, 19].

Fluid permeability describes fluid flow through a material in response to an applied pressure through the square of the limiting length scale to flow. One such limiting length scale could be the mesh size of EPS matrix filaments, which could be estimated using the scaling in Eq. 3,

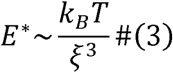

where *k*_B_ is the Boltzmann constant, *T* is temperature, and ξ is the correlation length or the mesh size of the network [69]. Using the calculated value for the EPS reduced elastic modulus (*E\** ), the mesh size, ξ, is on the order of 10’s of nanometers, resulting in an estimated permeability, *k* ∼ ξ^2^ ∼ 10^-16^ m^2^ (or 10^-10^ mm^2^ or 0.0001 µm^2^), on the same order of magnitude as estimated for high water content aqueous gels [70], including alginate gels (about 95% water) [71].

### 4.3 Viscoelasticity in Biofilms

Viscoelasticity is common in many biological systems, including biofilms [20, 72]. Fast viscoelastic responses protect cells within the biofilm from mechanical damage and facilitate the biofilm’s ability to stay intact under high pressures and instant forces [20, 27, 65]. Time scales of viscoelastic responses have been evaluated in soft, high-water content biological structures, like brain tissue, where viscoelastic contributions were found to dominate on time scales up to 100’s of seconds [73] (compared to slower diffusion-controlled structural relaxation, time scales ∼ 1,000’s of seconds [73–76]).

Stress relaxation data of pink berry granules were fit to viscoelastic models based on Maxwell elements, each consisting of two model components connected in series: a spring and a dashpot, representing elastic and viscous responses, respectively [74–76]. Fitting models were based on either one- or two-Maxwell elements. The two-Maxwell element model resulted in improved fits to the data (R^2^-values > 0.93 across all samples, N = 6) (Supplementary Information C and **Supplementary Fig. C1**). This two-element model has previously described the viscoelastic behavior of other multicomponent biological systems, including brain, breast cancer tissue, and meniscus extracellular matrix [73, 74]. Overall, our results indicate viscoelastic contributions to pink berry granule stress relaxation.

### 4.4 Limitations

In this study, in situ nanoindentation measurements enabled the selection of regions of interest. However, our samples are opaque unless sectioned into very thin slices. Slices thinner than 100 µm presented significant experimental and handling challenges due to their fragility. Therefore, nanoindentations were performed using 100 µm thick sections, despite the uncertainty in knowing where in the Z plane PSB microcolonies were positioned. In contrast, EPS regions were translucent and easily discernible with available optics. The uncertainty in spatial positioning of the nanoindenter with respect to microcolony distribution in the Z plane (normal to the sample holder) may explain the broad spread of *E\** measurements associated with PSB microcolonies and the relatively more consistent results for EPS matrix measurements (**Fig. 4** and **Supplementary Fig. B3**). In situ measurements with live cell fluorescent reporters and confocal microscopy may overcome these experimental challenges in the future.

It is important to note that the average reduced elastic modulus values for the EPS matrix and microcolonies in the pink berry were obtained from two centrally-located slices within the granule closest to the core, whereas the microindentation measurements were performed at the granule surface. Further studies are required to assess potential spatial variations in both density and stiffness of the components across the granule, as well as EPS composition to allow for comparative analyses with lamellar biofilms.While it would be ideal to conduct all imaging and mechanical analyses on the same pink berry samples, light sheet imaging requires fixation and optical clearing steps that would negatively impact subsequent mechanical characterization. For this reason, nanoindentations were not conducted on samples prepared for such imaging.

### 4.5 Future Work

The results presented here are a first approximation of the mechanical properties of pink berry granular biofilms. We believe there to be seasonal, geographical, and potentially day-night variability in properties like size and structure of the pink berries. Future studies will investigate variation in pink berry size, density, structure, and EPS matrix composition and explore how these variations impact mechanical properties of the biofilms. This work may help inform granule design and optimal environmental conditions in wastewater treatment systems.

Microscale variation in material properties throughout a granular biofilm can influence their macroscale properties and overall function. Radially symmetric density variation has been observed in other granular biofilms, typically with a denser surface and a less dense core [77]. While we did not find radially-symmetric trends from nanoindentations or imaging, a larger set of nanoindentation measurements positioned to investigate this property could improve our understanding of how spatial gradients influence overall mechanical properties and function.

EPS matrix variation, including composition and structured layering, have been found in other biofilms [78, 79]. Future studies will include investigation of EPS matrix composition and whether composition varies throughout the biofilms, as this could influence the mechanical properties of the composite.

Our results suggest that pink berry, soft and viscoelastic, pink berry granular biofilms are well suited for industrial applications. However, pink berries are only known to occur in a restricted geographic location worldwide and they cannot currently be maintained and cultured in isolation. This presents significant experimental challenges for detailed studies of component parts, or direct biotechnological applications. Future prospects include their propagation in culture or studies of live material using next-generation 3D culture and perfusion techniques [80].

Pink berry granular biofilms were used here as a model system for methods applicable across disciplines, integrating imaging and mechanics to construct structure-property relationships. We hope this study inspires future work on other biological composites and soft biomaterials and foresee a rich application space ranging from non-moldable synthetic materials to three-dimensional tissue-mimicking models like cancer spheroids and organoids.

## 5. Concluding Remarks

In this work, we have developed a sequence of methods bridging the gap between the visual and mechanical landscapes across length scales of composite soft materials. Using the soft (*E\**_pink berry_ ≈ 0.5–10 kPa) pink berry granular biofilm as a model system, we showed that this composite has viscoelastic mechanical properties and experience fast relaxation times (τ_1/2_ ≈ seconds). Additionally, our methods revealed that by computing the volume fraction of cell microcolonies (≈34%), we could relate microscale mechanical contributions to the macroscopic properties crucial for pink berry biofilm function, including structural integrity and cell protection. By studying naturally occurring microbial granules like pink berry biofilms, we can improve the design of engineered consortia, including for waste management technologies. Continued investigation of the structural and mechanical properties of biological composites, from biofilms to cancer spheroids, organoids, and beyond, will allow us to better understand nature and improve the built environment around us.

## Acknowledgements

We gratefully acknowledge the current and former members of the Interfacial Engineering Laboratory and the Wilbanks Lab for helpful discussions. Special thanks to Dr. Benjamin Lopez for his assistance with light sheet imaging and image analysis. This work was primarily supported by the US Army Research Office under Cooperative Agreement No. W911NF-19-2-0026 for the Institute for Collaborative Biotechnologies. We acknowledge the BioPACIFIC Materials Innovation Platform of the National Science Foundation (Award No. DMR-1933487) for the use of their shared user facilities, including the use of the Optics11 Life Pavone nanoindenter and instrument training and support from specialist Dr. Juan Manuel Urueña during nanoindentation measurements. We also acknowledge the use of the NRI-MCDB Microscopy Facility at UC Santa Barbara, including the Leica VT1000S vibratome, the Leica SP8 Resonant Scanning Confocal supported by the NSF MRI grant DBI-1625770 as well as the Zeiss Z.1 Lightsheet and Hercules computer workstation supported by the NIH Shared Instrumental Grant 1S10OD019969-01A1. We gratefully acknowledge contributions of Dr. Emily N. Junkins to field sampling of pink berry consortia, work which was supported by a Whitman Fellowship to E.N.J. from the Marine Biological Laboratory.

