## Supplemental Information for "Multiscale Mechanics of Granular Biofilms"

### A. Supplementary Figures – Whole Pink Berry Characterization

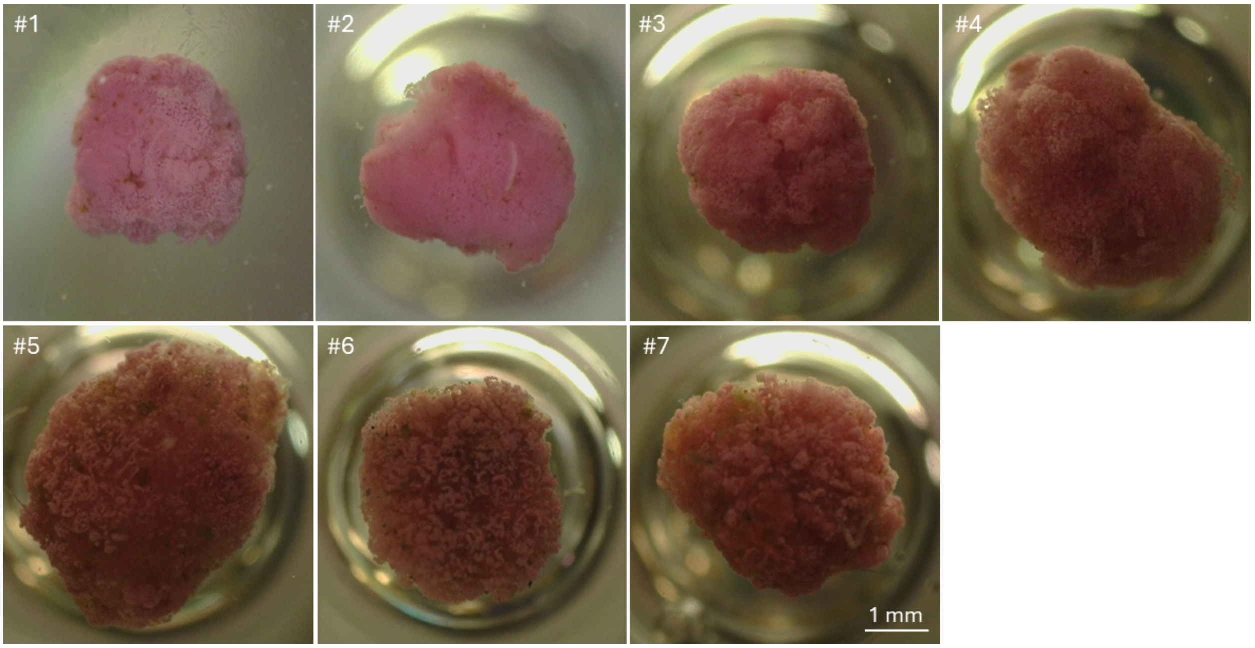


**Figure A1:** Images of pink berries used for microindentation measurements taken from below the indentation setup with the inverted digital microscope. Different shading of sample is from the probe casting a shadow from above on the sample, which is dependent on the z-position of the probe.

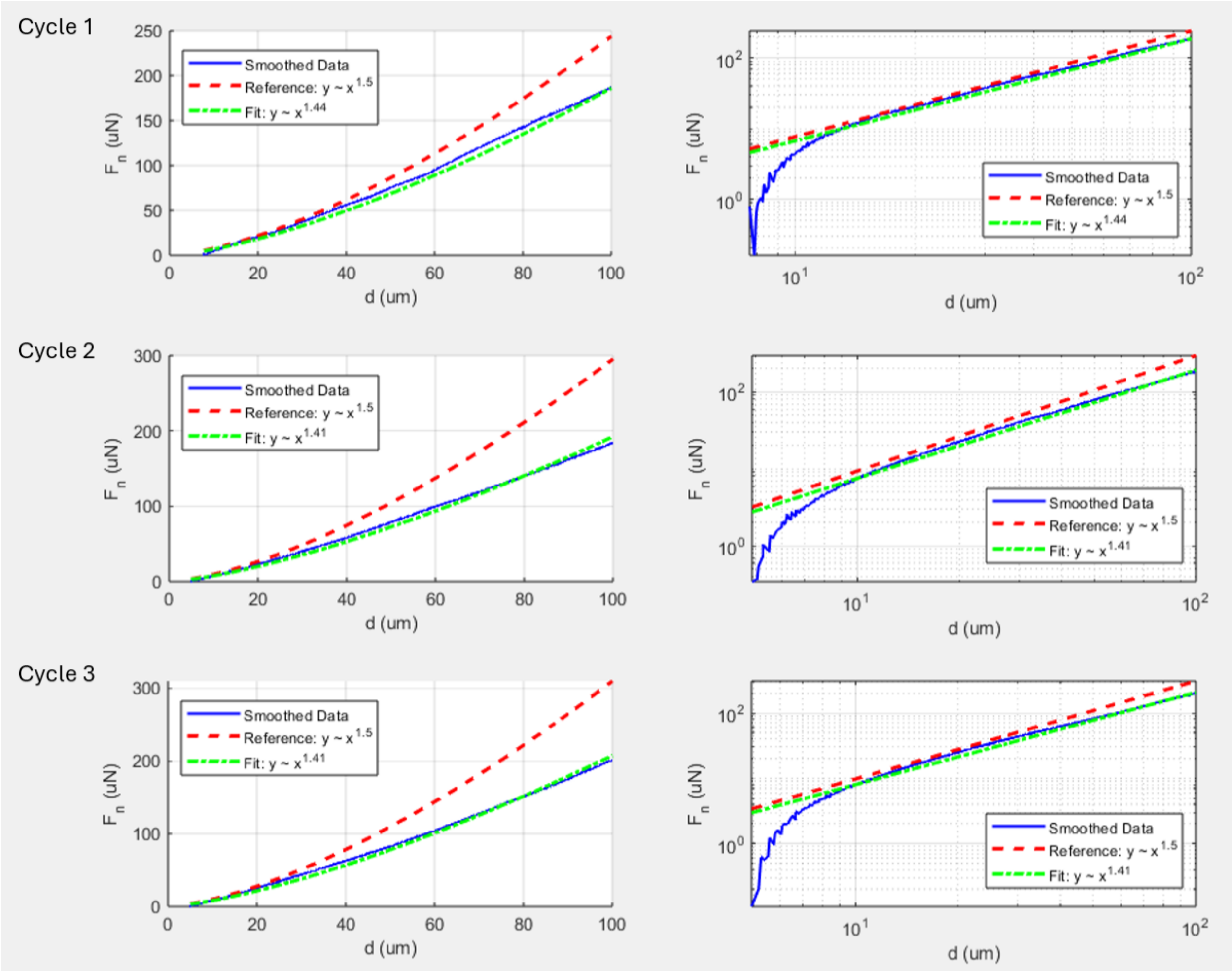


Figure A2: Force-displacement approach curves overlaid with $F{\sim}d^{3/2}$ scaling and best fit curves at low strains for three consecutive approach-retraction cycles of microindentation of a pink berry sample (N = 1). Right hand side plots are identical to the left-hand side data except plotted on log-log axes.

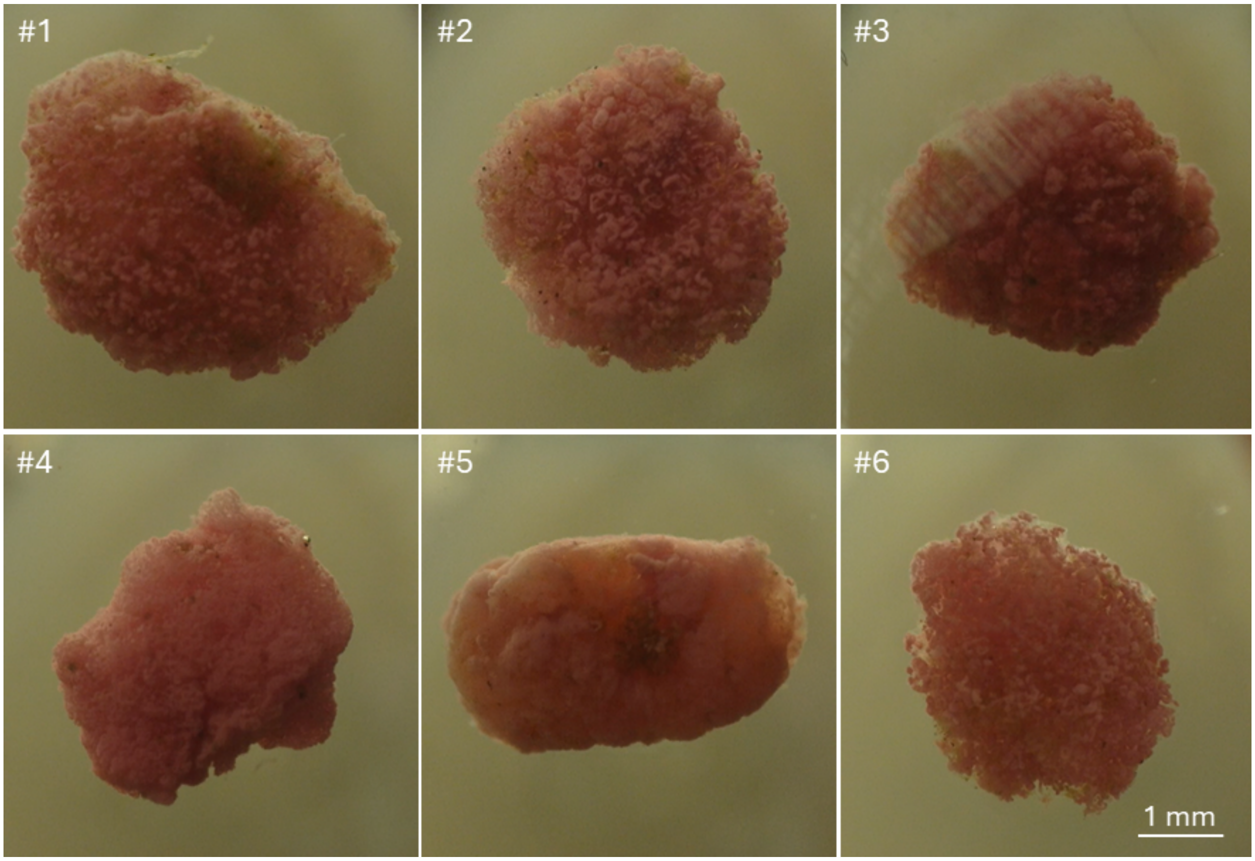


**Figure A3:** Images of pink berries used for stress relaxation measurements taken from below the indentation setup with the inverted digital microscope.

a) serial microindentation curves

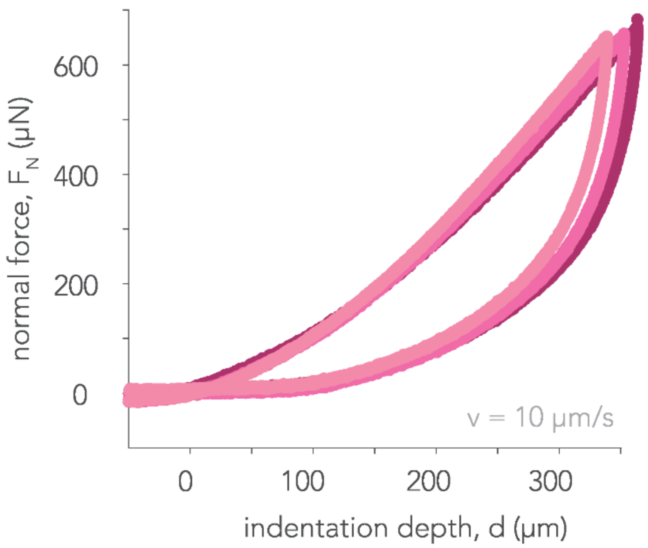


**Figure A4:** Serial indentation of pink berries shows very small, relatively unchanged, differences in mechanical behavior between cycles. For normal forces below 800 μN all pink berries examined show recovery after deformation from indentation, also tracked by video recording with a camera placed underneath the microindenter stage.

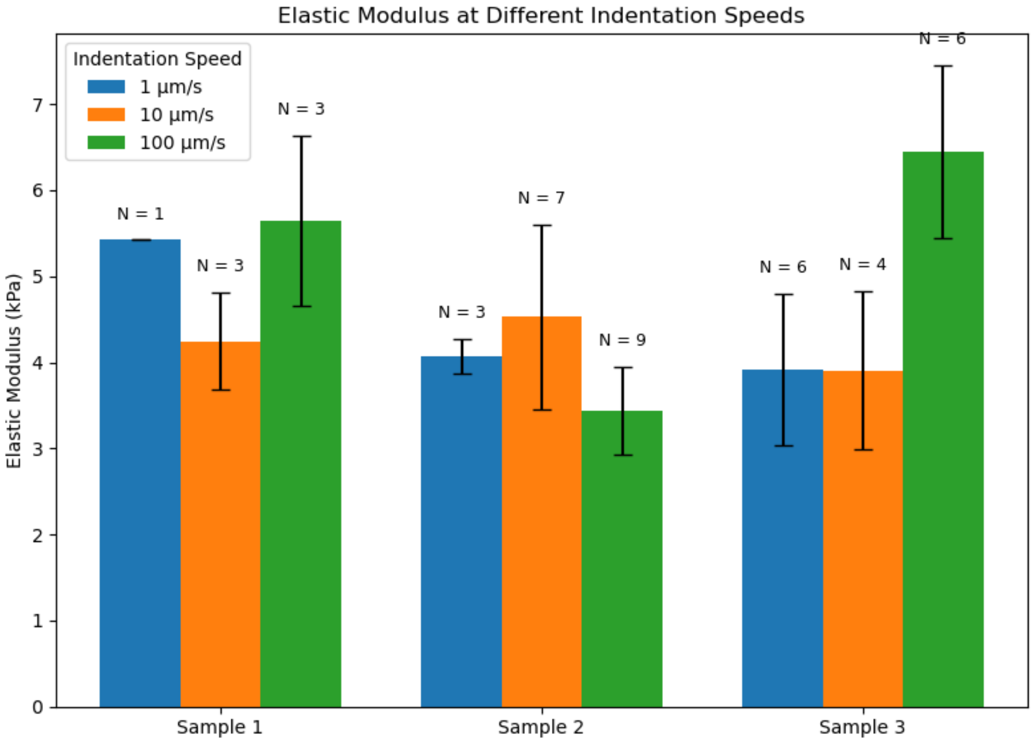


**Figure A5:** The reduced elastic modulus of pink berries is insensitive to the range of strain rates evaluated herein. Reduced elastic modulus values were calculated from fitting Hertzian contact mechanics models to force-displacement curves obtained over three indentation velocities (1, 10, and 100 µm/s) for three different pink berries. *N* indicates the number of approach-retraction cycles (technical repeats) for each velocity and each pink berry sample (labeled Sample 1, Sample 2, and Sample 3).

B. Supplementary Figures – Pink Berry Individual Component Characterization

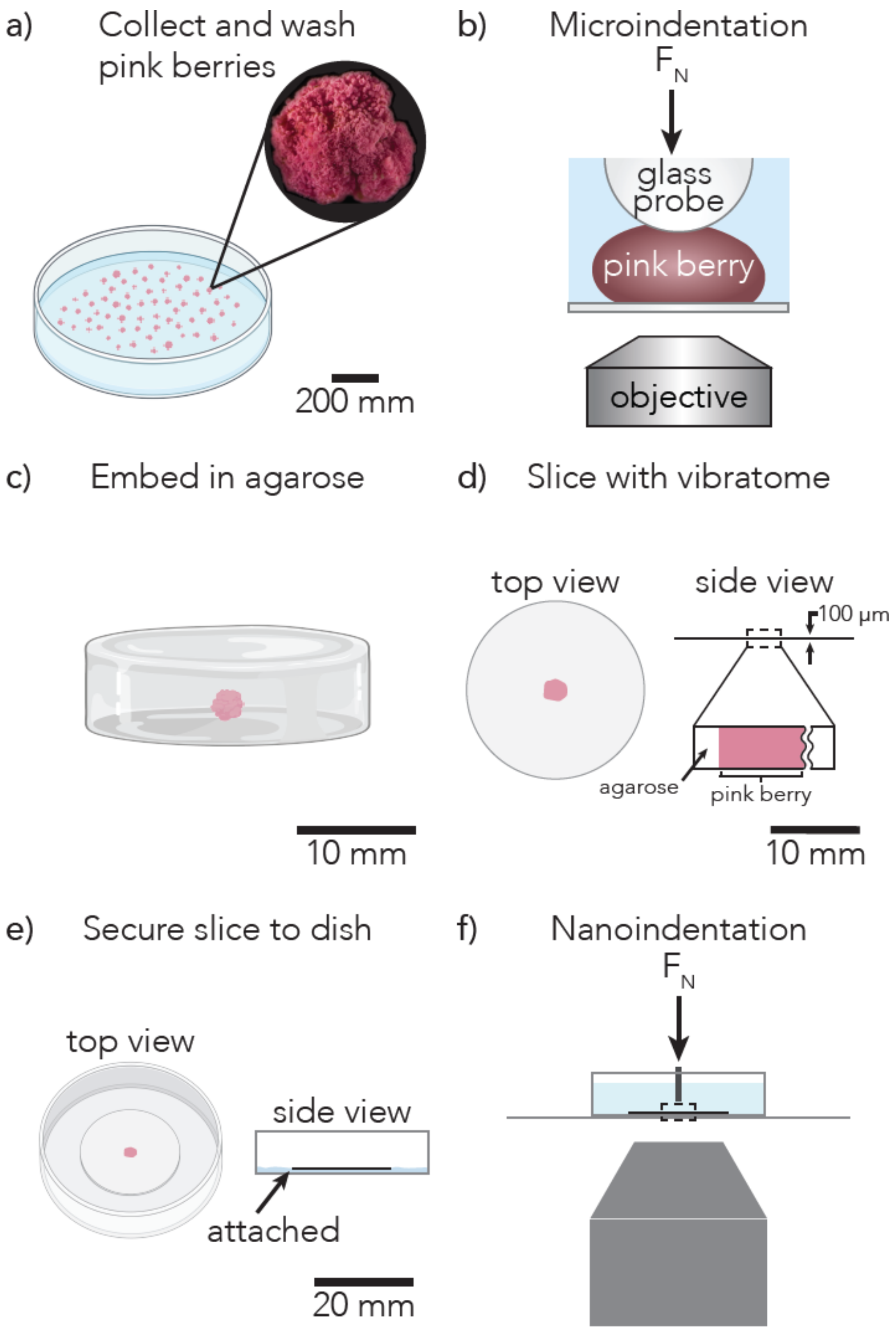


**Figure B1**: Overview of the methods presented for multiscale mechanical characterization of pink berry granular biofilms. Pink berries were collected and washed (a) before undergoing unconfined microindentation (b). Thereafter, the pink berry was cast in agarose gel (c) and sliced into ~100 micrometer thin sections on a vibratome (d). Using Weaver's gelatin coating on glass bottom petri dishes, the section was secured at the bottom of the dish (e) for nanoindentation measurements (f).

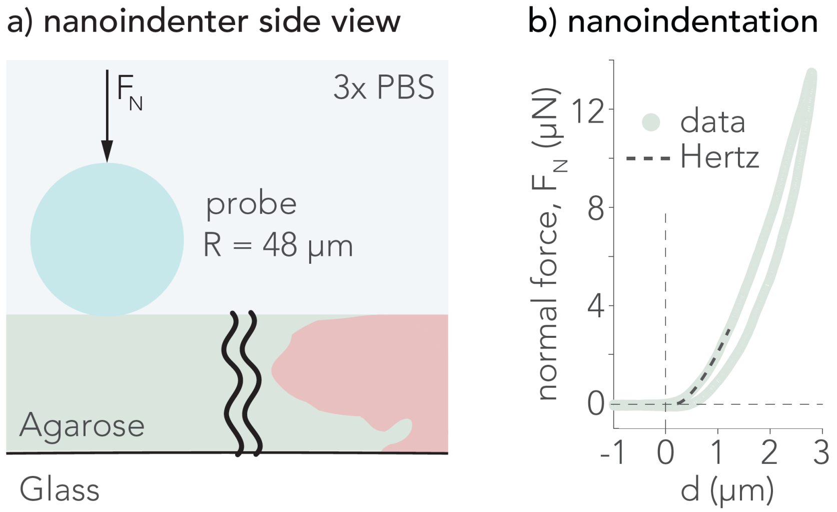


**Figure B2:** Agarose nanoindentation shows much larger reduced elastic modulus values compared to PSB microcolonies and EPS matrix in pink berry. a) Schematic side view of nanoindentation setup for agarose measurements. All measurements were taken with at least two probe diameters distance of separation, as well as away from the edge of the pink berry slice interfacing the agarose gel. b) Representative force-displacement curve from nanoindentation on agarose. Hertzian contact mechanics model was used to fit the approach curve for extraction of elastic modulus, shown as dashed lines on each curve. Reduced elastic modulus values for nanoindentations on 4 % (w/v) agarose gel (made in 3x PBS) ranged from 351 to 665 kPa, with a mean value of 471 kPa (N = 3 positions, standard deviation 139 kPa).

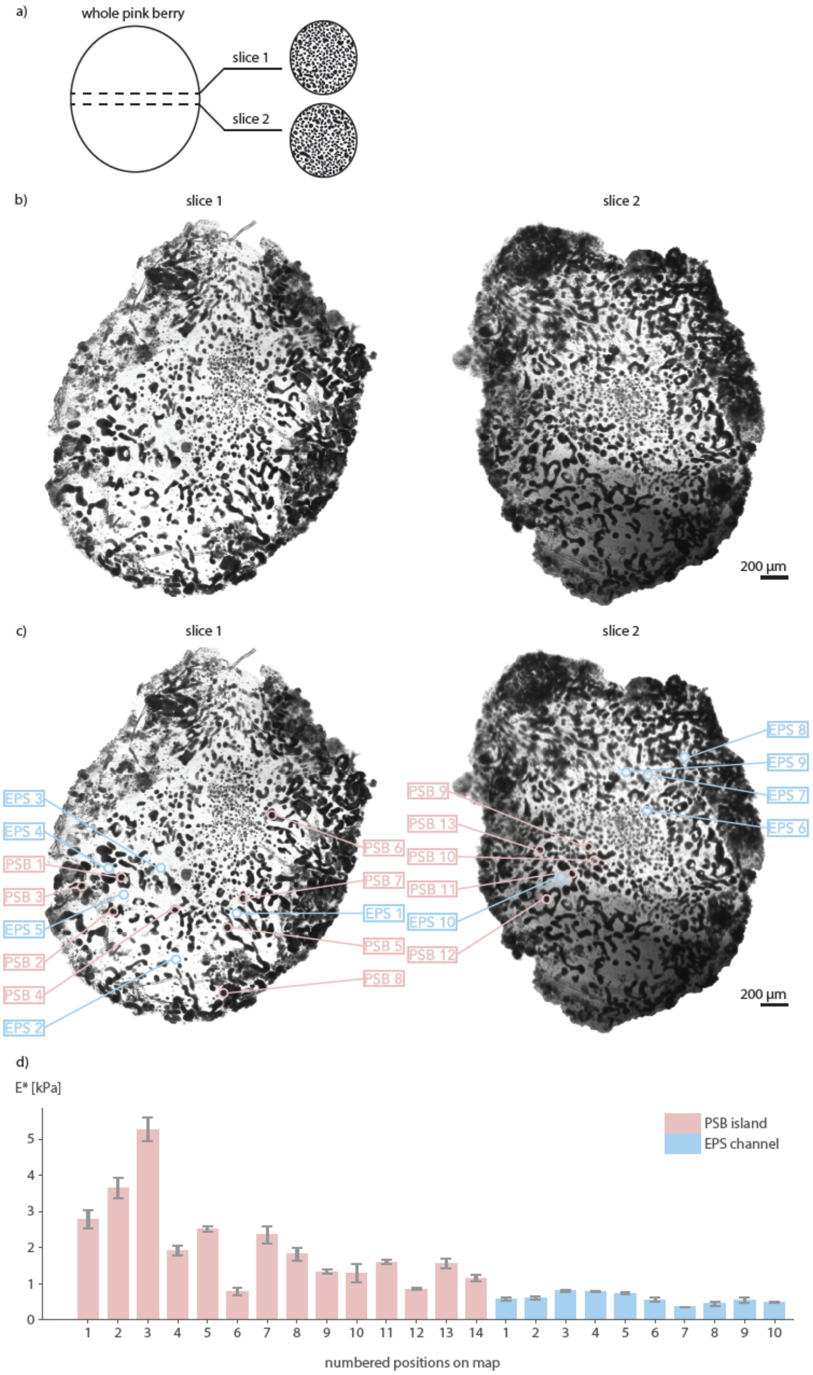


**Figure B3:** Overview of all positions indented with nanoindentation on pink berry sections together with their reduced elastic modulus values. a) Schematic of pink berry sectioning for nanoindentation measurements. Positions from two different slices from the same pink berry were indented. b) High resolution images of the two 100 micrometer thin sections prepared for nanoindentation measurements. c) Same as (b) but with all nanoindentation positions mapped out and enumerated in both sections. d) Reduced elastic modulus values per position enumerated as marked in (c). Barplots show mean values together with standard deviations in error bars from at least three indentations per position.

### C. Viscoelastic Model

Maxwell elements were used to fit normalized stress versus time from stress relaxation experiments to a viscoelastic model, where the following two equations were used. Pink berry samples measured show diverse enough stress relaxation to not fit one equation to all samples (see **Supplementary Fig. 10**), instead each sample has been fit individually with good $R^2$-values. Note that all stress relaxation data was normalized, therefore the *E* values below are not interpretable as they are unitless.

<u>1 Maxwell:</u>
A **single Maxwell element** (spring + dashpot in series) with a possible elastic part that never relaxes, $E_\infty$.

$$\frac{\sigma(t)}{\sigma_0} = E_1 e^{-\frac{t}{\tau_1}} + E_\infty$$

<u>2 Maxwell:</u>
**Two Maxwell elements** in parallel with a possible elastic part that never relaxes, $E_\infty$.

$$\frac{\sigma(t)}{\sigma_0} = E_1 e^{-\frac{t}{\tau_1}} + E_2 e^{-\frac{t}{\tau_2}} + E_\infty$$

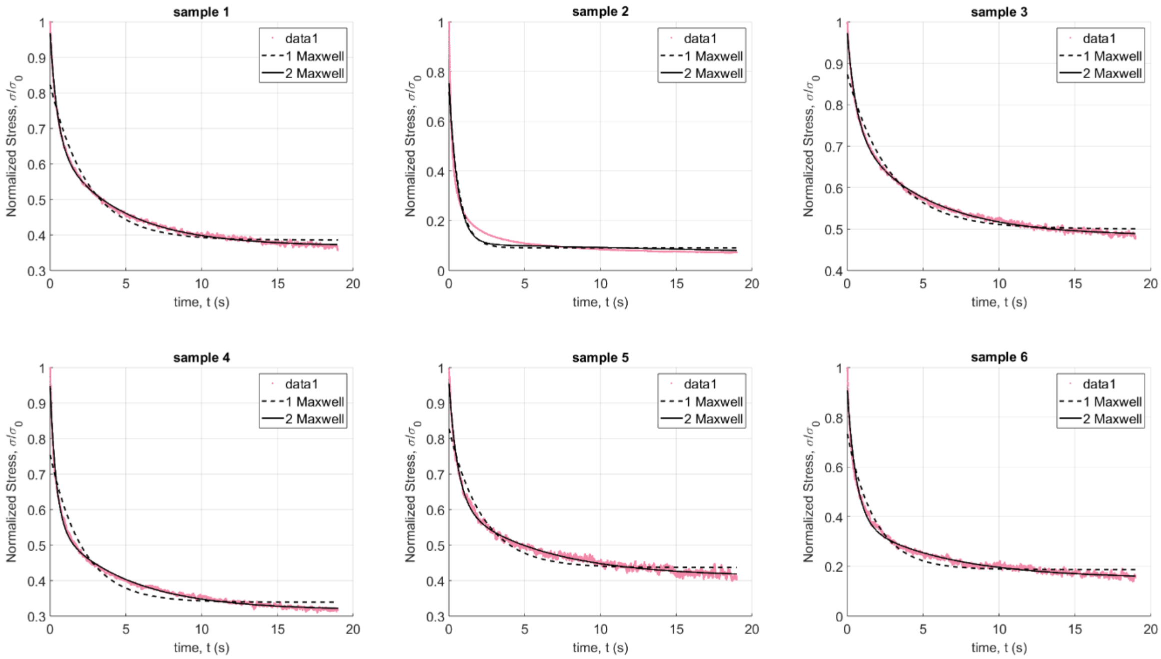


**Figure C1:** Stress relaxation data for all samples (N = 6) fitted to viscoelastic models with one and two Maxwell elements respectively. $R^2$-values are (from sample 1 to 6): 0.998, 0.939, 0.998, 0.996, 0.992, and 0.990.

Parameter fits for all pink berry samples (N = 6) fit to 1 Maxwell element

| Sample | $E_1$ | $\tau_1$ | $E_\infty$ | $R^2$ |
|---|---|---|---|---|
| 1 | 0.438378 | 2.46503 | 0.38546 | 0.965 |
| 2 | 0.630957 | 0.659328 | 0.0907276 | 0.913 |
| 3 | 0.372899 | 2.81941 | 0.5004 | 0.873 |
| 4 | 0.415294 | 2.12995 | 0.339036 | 0.935 |
| 5 | 0.391318 | 2.20417 | 0.436599 | 0.945 |
| 6 | 0.546014 | 1.81706 | 0.187196 | 0.945 |

Parameter fits for all pink berry samples (N = 6) fit to 2 Maxwell elements

| Sample | $E_1$ | $\tau_1$ | $E_2$ | $\tau_2$ | $E_\infty$ | $R^2$ |
|---|---|---|---|---|---|---|
| 1 | 0.314405 | 0.484248 | 0.287569 | 4.39111 | 0.368851 | 0.998 |
| 2 | 0.651189 | 0.566355 | 337524 | 2.55176e+08 | -337524 | 0.939 |
| 3 | 0.237254 | 0.611165 | 0.25362 | 4.9249 | 0.483769 | 0.998 |
| 4 | 0.391411 | 0.398887 | 0.241371 | 4.84669 | 0.317169 | 0.996 |
| 5 | 0.337678 | 0.584365 | 0.207835 | 5.80103 | 0.410703 | 0.992 |
| 6 | 0.52045 | 0.578893 | 0.239542 | 5.95298 | 0.151303 | 0.990 |

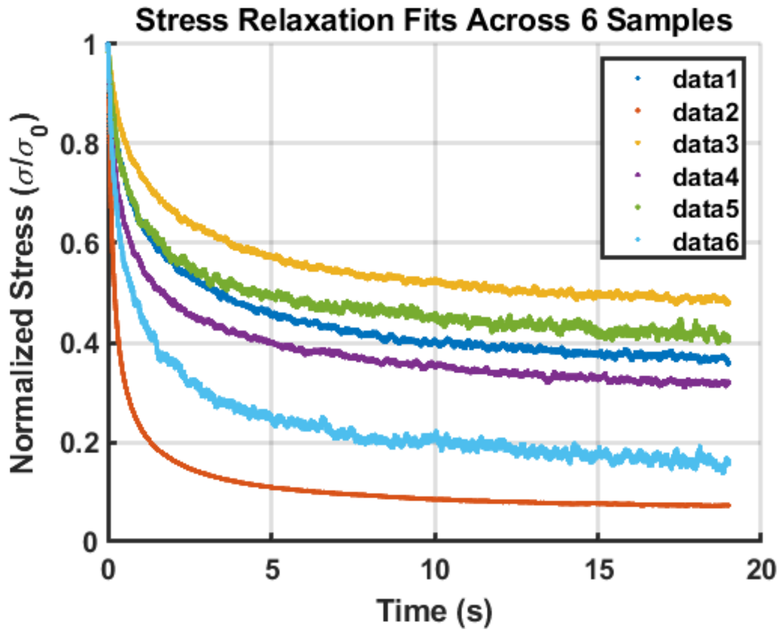


**Figure C2:** Normalized stress versus time data for all pink berry samples tested for stress relaxation (N=6).
